# Differential mutation accumulation in plant meristematic layers

**DOI:** 10.1101/2023.09.25.559363

**Authors:** Kirk R Amundson, Mohan Prem Anand Marimuthu, Oanh Nguyen, Konsam Sarika, Isabelle J DeMarco, Angelina Phan, Isabelle M Henry, Luca Comai

## Abstract

The upper plant body is formed by the continued growth of shoot apical meristems. In angiosperms, meristems are organized in three cell layers that tend to remain clonally isolated. Somatic variants emerge when mutant cells overtake part or all of a meristem. During sexual reproduction, only one layer, the L2, contributes to the next generation by forming gametes. The L2 is known to frequently invade and replace the L3, while L1-L2 separation is persistent. The role of different meristem layers in mutation accumulation is unknown. We discovered a potato periclinal chimera in which the L2 and L3, but not the L1, are marked by a chromosomal translocation. This enabled the identification of plants regenerated from leaf protoplasts originating from either the L1 or L2+L3. Leveraging these layer-specific clones, we identified and compared mutations that accumulated in the layers in the clonal parent for several decades. Here we show that the L1 accumulates mutations at 1.9 times the rate of the L2, indicating that plants might protect the germline by mechanisms that reduce the rate of mutation accumulation in the L2. In contrast to these layer-specific mutations, we found no evidence of somatic mutation fixation in all three meristem layers. Our findings highlight how periclinal chimeras are formed by independent mutational processes in which a mutation-prone epidermal layer could increase clonal variation.

## Introduction

Mutations fuel evolution but are often deleterious, and their rate is itself subject to selection ^1^. The arabidopsis mutation rate during sexual generations is 1-7 x 10^-9^ ^2^. Although the somatic mutation rate in differentiated tissues is as much as 1000 times higher ^3^, the number of fixed somatic mutations is low in most long-lived plants, and similar to the per-generation mutation rate in sexually reproducing plants ^4–9^. This suggests that plants have evolved mechanisms to control the mutation rate in stem cells.

The plant shoot stem cells reside in the shoot apical meristem (SAM) where they form a small cluster of undifferentiated initials from which all the body originates, including the germline ^10,11^. Slow growth of stem cells likely minimizes mutations by reducing cell cycles per sexual or clonal generation ^12^. In plants, the stem cells are organized in three clonally isolated histogenic layers called L1, L2 and L3 ^13,14^. In the meristem, L1 and L2 are single-cell layers in which cells divide only anticlinally. L1 generates the epidermis and trichomes, L2 part of leaf mesophyll, leaf veins, and gametes. The L3 layer is made of cells that divide both anticlinally and periclinally, forming a tridimensional ensemble that generates the vascular system, ground tissue, and in vegetatively propagated plants, the roots.

The layers maintain separation in the SAM, although occasionally cells from one layer invade another. In axillary meristems of potato and arabidopsis, cells of the L2 often invade the L3 ^14,15^. Separation of the L1 from the L2, however, is more persistent, as demonstrated by the existence of stable periclinal chimeras in which L1 is mutant compared to the L2/L3. For example, Pinot Meunier, a clonal variant of Pinot Noir, is a chimera in which the L1 over-accumulates the hormone gibberellin, while L2 and L3 are wild-type and identical to Pinot Noir ^16^. During tissue culture of Pinot Meunier, the two cytotypes can arise separately as uniform plants, presumably by regeneration of a single L1 or L2 or L3 cell. Other vegetatively propagated crops, such as potato and citrus, regularly undergo somatic mutation and generate new types called sports that are often agronomically valuable ^17^.

Despite its basic and economic importance, the relation between mutation rate, meristem layers, clonal and sexual variants is poorly understood. In peach, excision of a transposable element inserted in an anthocyanin biosynthetic gene was four times more frequent in the L1 than in the L2 ^18^. In arabidopsis, genome protection pathways are preferentially expressed in L2 meristem cells ^19^. This raises the question of whether protection mechanisms are in place to reduce mutation accumulation in the L2, thus sheltering the germline.

The modern varieties of potato are clonally propagated, highly heterozygous autotetraploids. During tissue culture regeneration of protoplasts, a procedure required for transgene-free editing, potatoes display frequent phenotypic changes in a phenomenon called somaclonal variation ^20^. In the variety Red Polenta, we have demonstrated that regeneration is associated with genomic instability manifested as aneuploidy and chromosomal aberration ^21^. Here, we further characterized these variants. We discovered a preexisting chromosomal translocation that was present in Red Polenta L2 and L3 cells, while the L1 retained the ancestral karyotype. Red Polenta is, therefore, a periclinal chimera. Protoplast regeneration produced pure types in which presence or absence of the translocation marked the layer of origin. This allowed comparison of how fixed somatic mutations accumulated in L1 and L2 since Red Polenta diverged from its clonal parent, cv. Urgenta. We show here that mutations accumulated differentially in the two layers and that they are more frequent in the L1. Differential sensitivity to natural mutagenesis may help explain the success of the layered meristems of angiosperms.

## Results

In a previous study, plants were regenerated from individual leaf protoplasts of tetraploid potato cultivar Red Polenta ^21^. To better characterize the changes that might have occurred during the regeneration process, we characterized the genomes of twelve of these plants in detail. Each regenerated plant was sequenced to ≥10x read depth per haplotype with short reads (Supplemental Table S1).

### Red polenta is a periclinal chimera

As described previously ^22,23^, Red Polenta is a clonal variant of cv. Urgenta but distinguishable by a chromosome translocation called tr8-7 (three copies of chromosome 8 4.6 Mb distal right arm, five copies of chromosome 7 5.6Mb distal right arm). Surprisingly, six of the 12 lines independently regenerated from Red Polenta lacked the segmental aneuploidy typical of tr8-7. Short read depth analysis demonstrated the loss of CNVs associated with tr8-7 (Fig. 1A, Supplemental Fig. S1). As a result of regeneration, the six plants appeared to have transitioned from an aneuploid to a euploid state with respect to this chromosome region. In the six euploid regenerants, the region of chromosome 8 involved in the translocation displayed 13,542 SNP in simplex dosage (1 variant : 3 reference allelic ratio). These SNPs were grouped in distinct clusters along the end of chromosome 8. They were not detected in Red Polenta. The SNPs were also absent in the regenerants that showed tr8-7 and from Red Polenta roots (Fig. 2B).

**Figure 1.**
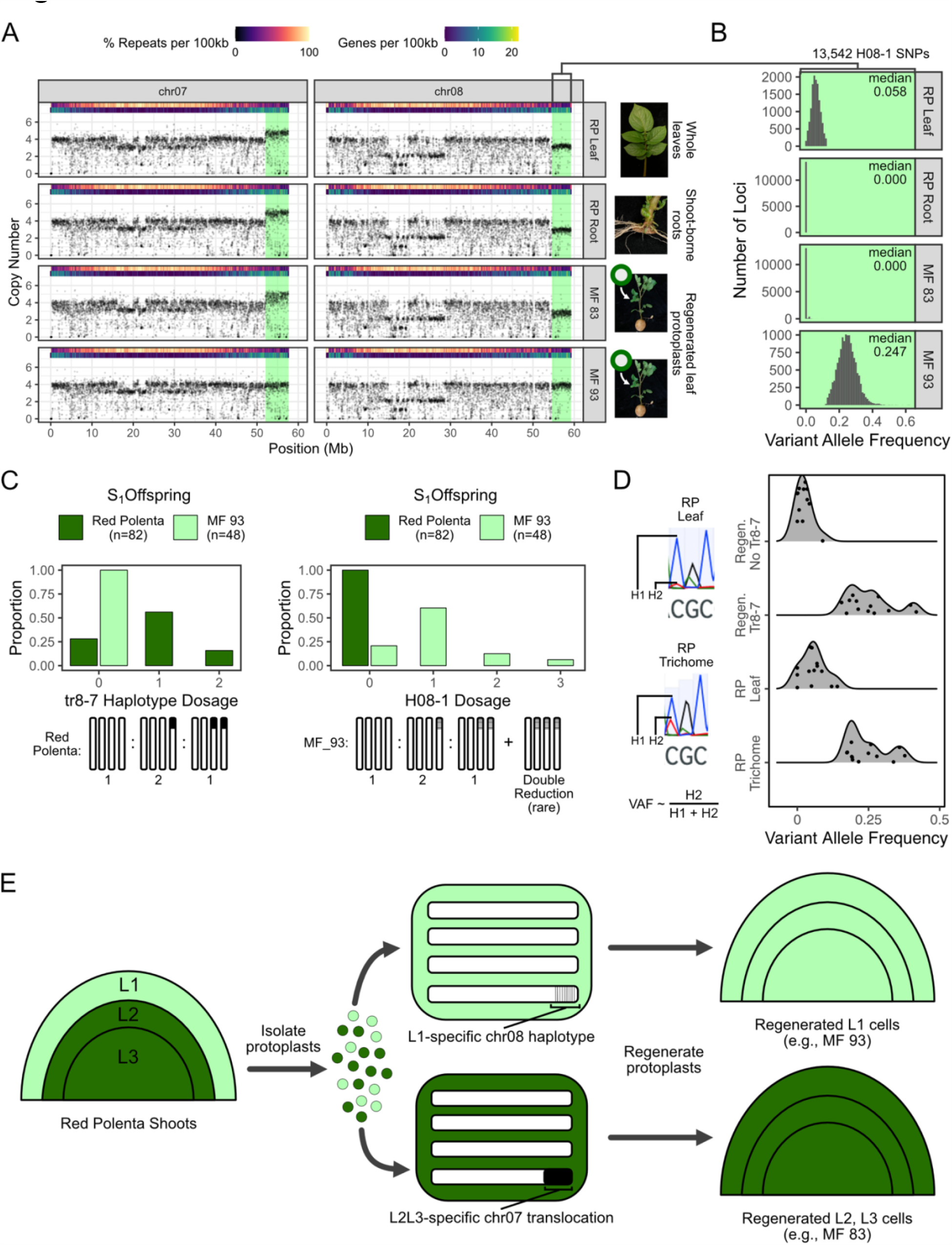
Periclinal chimerism in Red Polenta. **A)** Read coverage plots display five copies of the tip of chromosome 7 and 3 copies of the tip of chromosome 8, indicating the presence of the translocation tr8-7 in leaf and root cells of Red Polenta (RP). The same signal is present in regenerant MF83 but absent in regenerant MF93. **B)** Variant Allele Frequency (VAF) for 13,542 SNPs spanning the tip of chromosome 8. This haplotype, called H08-1, is substoichiometric in Red Polenta leaves because it is only present in the L1 layer. It is absent in Red Polenta roots because they are formed preponderantly from the L3. It is also absent in L2+L3 regenerant MF83 and present at the expected simplex VAF in L1 regenerant MF93. **C)** Inheritance of tr8-7 and haplotype H08-1. Self-fertilized offspring of Red Polenta (n=82) exhibited Mendelian segregation with respect to tr8-7 but complete absence of haplotype H08-1. On the other hand, MF93 and its offsprings completely lacked tr8-7 but haplotype H08-1 segregated as expected, including a few cases of double reduction **D)** Selected SNPs from haplotype H08-1 were PCR amplified and scored by Sanger DNA sequencing. The VAF illustrates the normal simplex state in an L1 regenerant, its absence in an L2+L3 regenerant, the low VAF resulting from minority state in Red Polenta leaves, and the VAF corresponding to simplex allele dosage in Red Polenta trichomes. **E)** Model of chimerism and isolation of clonal derivatives. Red Polenta is a periclinal chimera in which the L1 layer is wild-type with respect to tr8-7 and has a unique, ancestral chromosome 8 haplotype. The L2 and L3 layers are mutant and share translocation tr8-7.

**Figure 2.**
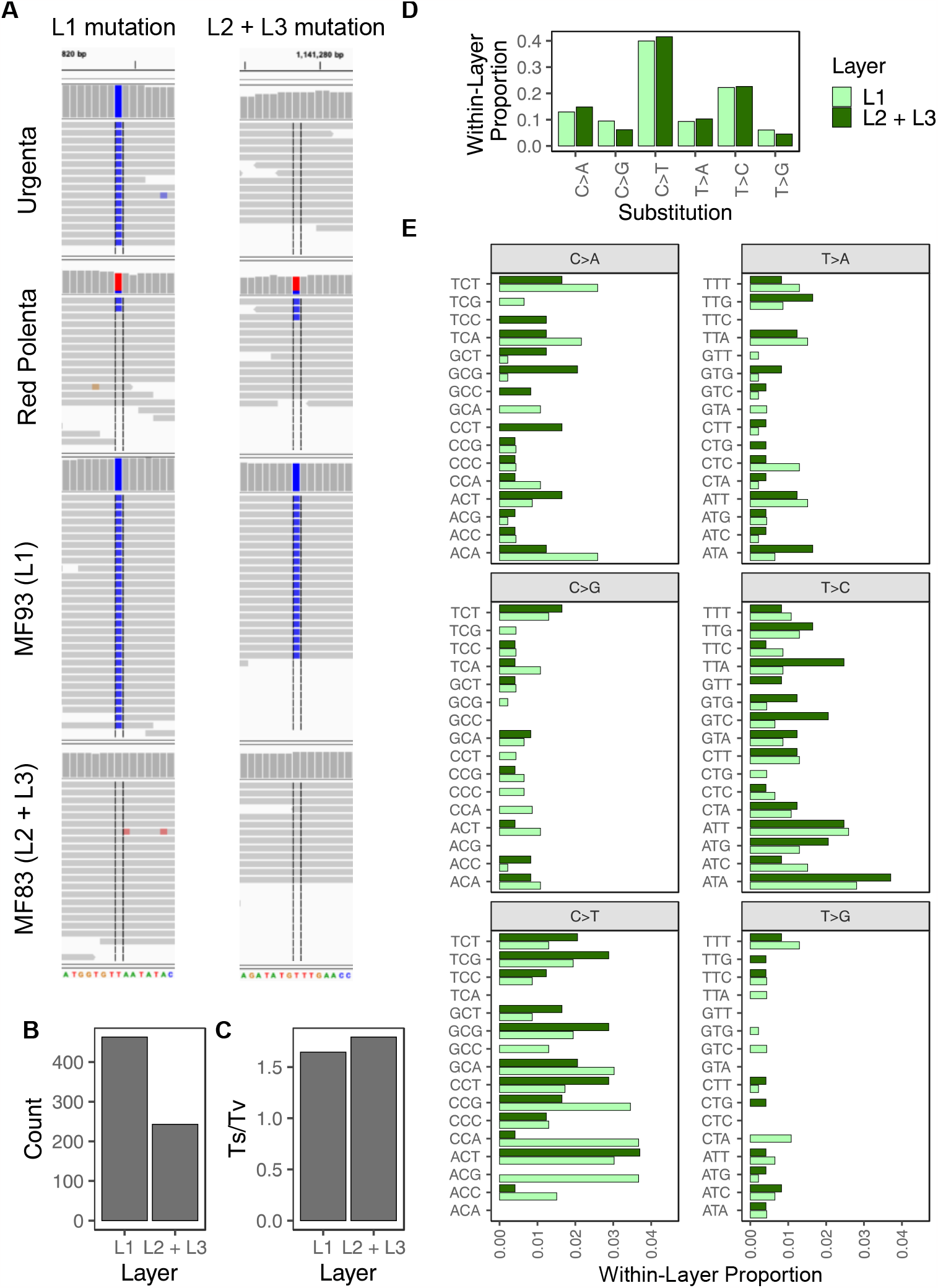
Layer specific mutations. **A)** Example of single nucleotide variants specific to the L1 or L2+L3 layers. Sequence reads from four genotypes were mapped to specific, single haplotype assembled from Red Polenta leaf DNA. Because the unitig represents a unique haplotype, a fully reference or a fully variant signal is expected from regenerated plants. In this system, Urgenta is assumed to carry the ancestral allele. At the locus displaying an L1-specific mutation, Urgenta and regenerated L2+L3 are fixed for the same allele, while regenerated L1 is fixed for the other allele. In Red Polenta, the chimeric state results in approximately 20% VAF of the L1-specific allele. At the locus displaying an L2+L3-specific mutation, Urgenta and regenerated L1 are fixed for the same allele, while regenerated L2+L3 is fixed for the other allele. Both alleles appear in Red Polenta, with the L2+L3-specific mutant allele represented by approximately 80% of reads. **B)** The total count of unitig-mapped mutations. L1 mutations are 1.9X more frequent than L2 mutations over the 1.05 Gb assayed. **C)** Relative frequencies of transitions and transversions in cell layers. **D)** Relative frequencies of each of the six single-nucleotide substitution types in cell layers. Differences in any one substitution type were not significantly different between layers. **E)** The trinucleotide spectra of L1 and L2+L3 mutations display overall similarity with differences in specific contexts.

A preliminary hypothesis posited that these SNP arose from error prone replication after mitotic recombination between homologous chromosomes, such as during Break Induced Replication ^24^. Further characterization, however, ruled this out because the SNPs were common among potato cultivars and including cultivars that predate Urgenta, such as Irish Cobbler, Garnet Chili and Early Rose, as well as landraces and wild species with no recorded ancestry in the pedigree of Urgenta (Supplemental Fig. S2). A reexamination of leaf DNA from Red Polenta revealed that these SNP were in fact present in Red Polenta at low variant allele frequency (VAF, median: 5.8%) but had been previously filtered out for deviating from the 25% VAF expected for simplex genotypes (Fig. 1B). To reconcile these unusual observations, we hypothesized that Red Polenta is a periclinal chimera in which most cells of the shoot carry tr8-7, while a small proportion of cells retain the ancestral chromosome 8 haplotype, including the 13,542 newly uncovered SNPs. This could be explained if this proportion of cells represented a single meristem layer of Red Polenta.

### The L1 layer carries the ancestral allele

We next set out to determine which Red Polenta meristematic layer carries the mutant allele. Cytological detection of tr8-7 in root cells ^22^, combined with the near absence of putatively layer-specific chromosome 8 SNPs indicated that the L3 layer was mutant (Fig. 1A, 1B). Because gametes are produced from the L2 layer, we examined inheritance of tr8-7 in Red Polenta selfed progeny. As predicted for tetrasomic segregation of a simplex allele, the progeny displayed 1:2:1 inheritance for 0, 1 or 2 copies of tr8-7 (Fig. 1C). This confirmed tr8-7 the previously observed meiotic transmission observed when Red Polenta was used for dihaploid induction ^22^. We also investigated the transmission of the chromosome 8 SNP set. All Red Polenta progeny lacked the set, while it was inherited according to tetraploid simplex genetics among offspring of euploid regenerant MF93, with three exceptions due to double reduction (Fig. 1D, Supplemental Fig. S3). These results are consistent with exclusion of the chromosome 8 SNP set from the Red Polenta germline due to confinement in L1 rather than negative selection in the germline. Finally, to investigate the state of the L1, we extracted genomic DNA from Red Polenta trichomes and Sanger sequenced PCR amplicons covering 13 of the chr08 SNPs with predicted L1-specific alleles. In the chromatogram peaks, we measured the ratio of the predicted L1-specific allele to the total signal (Fig 2E), and observed a mean of ∼0.25 for both the L1 regenerant and the trichome DNA. The peak was near zero in the regenerants with tr8-7 (from L2 + L3), and ∼0.05 in leaf DNA of Red Polenta. Taken together, these data indicate that the L1 of Red Polenta carries an ancestral, euploid karyotype, whereas L2 and L3 carry the tr8-7 mutation (Fig. 1F).

Periclinal chimeras can display considerable stability over long periods of vegetative propagation ^25^. To investigate the stability of the Red Polenta periclinal arrangement, we reanalyzed short read sequencing of a Red Polenta holding that has been propagated independently for at least the last 25 years, and found that it is also a tr8-7 periclinal chimera (Supplemental Fig. S4). Based on chimeric layer behavior ^14,15,26^, we speculate that the mutation arose in L2 and invaded L3. Stability of a chimeric SAM is surprising in this case because it involves aneuploidy, which is typically deleterious even when buffered by polyploidy.

### L1 stem cells carry almost twice as many mutations as L2/L3 stem cells

Regeneration from single cells of Red Polenta yielded homogeneous plants of one or the other type. The ability to isolate single, genetically marked cells from the different meristem layers provided a unique opportunity to examine the mutation rates and spectra of cell lineages in the Red Polenta SAM.

Potatoes are highly polymorphic and heterozygous, making accurate and robust mutation detection challenging, especially when attempting to detect layer-specific mutations. To facilitate mutation discovery, we obtained 15x per haplotype long reads from Red Polenta and assembled a draft genome assembly consisting of haplotype-resolved unitigs (unitig N50=349 kb). We next mapped Illumina reads to this Red Polenta assembly and identified single-nucleotide mutations (see Methods). We excluded likely false-positive variants near homopolymers and limited our analysis to the contigs that represented single haplotypes (haplotigs). This represented approximately 1.05 Gb of assayable bases, amounting to 71% of haplotig space and 36% of the unitig assembly. The 13,542 simplex SNP set in the chromosome 8 L1-specific haplotype provided a template for mutation discovery: mutations should appear as polymorphic sites that are unique to a regenerant class. When uniquely mapping reads to a single-copy haplotype, variants are expected to be fixed for different alleles in the two regenerant classes, with the L1-specific allele appearing at substoichiometric VAF in Red Polenta (Fig. 2A). To infer the ancestral and derived alleles, we analyzed whole-genome sequencing of an Urgenta holding from the USDA ^23^; we assumed it carried the ancestral allele. Using this method, we identified 463 de novo mutations that were unique to the L1 layer, and 243 mutations that were unique to the L2+L3 layers. This represents a 1.9-fold difference in somatic mutation accumulation between cell layers (Fig. 2B).

In the SAM, L1 and L2 are typically maintained as clonally distinct layers due to cell divisions primarily occurring in the anticlinal plane. This property tends to inhibit somatic mutations arising in any one layer from becoming fixed throughout the entire meristem. We next leveraged our regenerants to see if we could detect such mutations present in all three meristem layers on the 1.05 Gb of Red Polenta haplotigs. We considered loci for which Red Polenta and all regenerants were fixed for one allele, and two Urgenta holdings, one from the USDA, the other from IPK (Gatersleben, Germany), were both fixed for the same alternate allele. No such loci were found, confirming that the stratified SAM strongly inhibits fixation of somatic mutations throughout all three Red Polenta meristem layers.

To investigate processes that may have driven the difference in somatic mutation accumulation between cell layers, we compared the mutation spectra of the L1 and L2+L3 layers. The transition:transversion ratio was not significantly different between L1 and L2+L3 (*p*=0.66, χ^2^ test) (Fig. 2C). Next, we considered the six possible single-nucleotide substitutions: for example, C-to-T and G-to-A are equivalent mutations located on opposite strands. We found that C-to-T transitions were the most common type of substitution in both L1 and L2+L3 (Fig. 2D). However, the proportions of any one substitution type were not significantly different between layers (χ^2^ test). Mutation trinucleotide context can predict biological processes and environmental exposures. To identify an underlying mutagen, we used SigProfilerExtractor ^27^, a nonnegative matrix factorization method, to compare L1 and L2+L3 mutation spectra to the COSMIC mutation database ^28^. Analysis of L1 mutations identified one mutation signature that represented a mixture of two COSMIC mutation signatures: SBS25 and SBS5. Reconstruction of these signatures produced a cosine similarity of 0.693 with the observed L1 mutations (Supplemental Fig. S5A). Similarly, analysis of L2+L3 mutations identified a one mutation signature representing a mixture of SBS5 and SBS87, and reconstruction of these signatures produced 0.644 cosine similarity with the observed mutations (Supplemental Fig. S5B). Taken together, our analyses reveal differences in the rate, but not the spectrum, of single-nucleotide somatic mutation accumulation between histogenic layers.

## Discussion

Our analysis of the Red Polenta system reveals that the L1 cell layer accumulates nearly twice as many somatic mutations as the combined L2 and L3 cell layers. At least four mechanisms might explain this observation. i) faster cell division ii) higher exposure to environmental mutagens, iii) relaxed epigenetic of TEs and connected error prone repair of dsDNA breaks, and iv) inefficient DNA repair. The first explanation is inconsistent with experimental findings ^29^. Regarding the second hypothesis, developing leaves shield the SAM from mutagenic radiation, although oxygen and other gasses are likely to form a gradient from surface to internal cells ^30^. To test this hypothesis, we compared the transition:transversion ratio and frequencies of substitution types of layer-specific mutations, and did not observe significant differences between layers. Further comparison of trinucleotide context of layer-specific mutations identified similarities to known mutational signatures in the COSMIC database, but the relevance of human mutational signatures to potato should be kept in mind when interpreting these results.

The last two explanations are both consistent with the limited data available in the literature ^18,19,31^. Increased mutation accumulation in the L1 expands our understanding of the layered meristem function. Seedless vascular plants typically exhibit a point meristem with a single initial cell, while many gymnosperms and most angiosperms have developed stratified meristems. Did this innovation contribute to their explosive radiation? A layered meristem facilitates the persistence of deleterious mutations ^32^. Dosage sensitive mutations, on the other hand, may change epidermal properties providing tolerance to a pest or environmental stress, or even affect development of other layers ^33–35^. An increased rate of mutation accumulation in the L1 may enable the testing of innovations relevant to the epidermis without incurring a germinal penalty. If an L1 mutation provides a selective advantage, invasion of L1 into the L2 could provide an avenue for genetic fixation. Studies of periclinal chimeras indicate that such invasions do occur ^36,37^, however, we did not find genetic evidence of invasions leading to somatic mutation fixation in Red Polenta meristems.

## Methods

### Plant materials and sequencing

Cuttings were propagated *in vitro* under 16h light 25°C: 8h dark 18°C on half-strength Murashige and Skoog medium adjusted to pH 5.7 with KOH and supplemented with 1% sucrose, 1x Gamborg vitamins and 0.5g/L MES. Single-node cuttings were transferred to fresh media every 1-2 months. Red Polenta and protoplast-regenerated plants were continuously propagated by tubers on soil in 8” pots in ambient greenhouse conditions at Davis, CA, USA. The topmost three leaves of five Red Polenta plants grown from tubers of the same mother plant were collected for high molecular weight DNA isolation and library preparation. Seeds produced from self-pollination of MF93 were soaked in 2000 ppm GA3 for 24h, rinsed thoroughly with sterile water, surface sterilized in 50% commercial bleach for 5 minutes, and rinsed an additional 5 times with sterile water. Seeds were then germinated on half-strength Murashige and Skoog medium in the aforementioned growth chamber conditions. Cuttings were propagated every 1-2 months as described above.

High molecular weight DNA was isolated from 1.5 grams of Red Polenta leaves using a sorbitol pre-wash and high-salt CTAB extraction as previously described ^38^. Quality was assessed on a Bioanalyzer, and a HiFi library was then prepared according to the manual “Procedure & Checklist -preparing HiFi SMRTbell Libraries using SMRTbell Express Template Prep Kit 2.0”, including initial DNA fragmentation by g-Tubes (Covaris) and final library size binning by SageELF (Sage Science, Beverly, MA). Size distribution was controlled using a Bioanalyzer again before sequencing. Size selected libraries were sequenced on a Sequel IIe instrument at the University of California Davis DNA Technologies Core (Davis, CA, USA) with Binding kit 2.0 and Sequel II Sequencing Kit 2.0. For short read sequencing, genomic DNA was extracted from greenhouse or *in vitro* grown plants and used for library preparation as previously described ^39^. All libraries were prepared and sequenced as specified in Supplemental Table S1. Sequencing reads from previous studies ^21–23^ were retrieved from the National Center for Biotechnology Information (NCBI) Sequence Read Archive (SRA) and incorporated in subsequent analyses.

### Sequence processing and alignment to DM1-3 reference genome

Short reads were trimmed with Cutadapt 1.15 ^40^ and aligned to the DM1-3 version 6.1 assembly with BWA mem (version 0.7.12-r1039) ^41^. PCR duplicates were removed with Picard (version 2.18) MarkDuplicates. For all paired-end read alignments with overlapping mates, the overlap region was soft-clipped for one of the two mates using bamutil:clipOverlap (version 1.0.14) ^42^. Paired ends with mates aligning to different contigs were removed. These processed alignments were used for read depth and variant calling analyses.

### Layer-specific variant discovery and genotyping

#### Variant discovery

Copy number was inferred from median read depth of non-overlapping 10kb windows of the DM1-3 reference genome ^43^ as previously described ^23^. Upon observing polymorphism with respect to tr8-7, we assayed all regenerants for single-nucleotide and short indel mutations. Variants were called and genotyped using freebayes (version 1.3.4) ^44^ with minimum mapping quality 40, minimum base quality 20, population priors off, a maximum of six alleles considered per variant and all other parameters left at the default setting. To remove low-quality variants, the following hard filters were applied using bcftools (version 1.15) ^45^: TYPE=‘snp’, NUMALT=1, EPP≤30, EPPR≤30, MQM≥30, MQMR≥30, RPP≤30, RPPR≤30, SAP≤30, SRP≤30. Only sites with Red Polenta read depth between 40 and 140 were retained. Novel variants associated with the presence of tr8-7 were defined as ≥12.5% VAF in each of the six regenerated lines that lacked tr8-7 and ≤12.5% VAF in each of the six regenerated lines that exhibited tr8-7. The 13,542 such SNPs located to the right of chr08:54,585,000 were defined as the H08-1 haplotype.

#### Diversity panel variant genotyping

Publicly available short read sequencing of 102 partially redundant samples from previous studies ^22,23,39,46^ was retrieved from NCBI SRA for processing and alignment as described above. Processed alignments were used to genotype each of the 102 samples at the 13,542 H08-1 SNP loci with freebayes version 1.3.4. We applied filters to retain only biallelic SNPs (NUMALT=1 & TYPE=‘snp’) with ≤10% missing data to the resulting VCF, which retained 5,980 H08-1 loci. All samples corresponding to Red Polenta and other duplicates were removed. For example, multiple Desiree holdings were included for variant calling, but only one was retained. This yielded a nonredundant panel of 76 samples. Allele frequencies were recalculated to reflect the nonredundant samples before plotting the allele frequency spectrum (Supplemental Fig.S2).

#### Offspring genotyping

To assay tr8-7 dosage of 82 Red Polenta S1 offspring, chromosome dosage analyses were carried out as previously described ^21,47^. Standardized coverage values for each offspring were normalized to Red Polenta and multiplied by four to indicate a tetraploid state. No correction was made for PI310467 copy number. Dosage genotypes were derived from the average standardized coverage values of the dosage-variable ends of chromosomes 7 and 8. For chromosome 7, these genotypes were “down7” if the average standardized coverage was less than 3.9, “mid7” for coverage between 3.9 and 4.75, and “up7” for exceeding 4.75. For chromosome 8, the genotypes were “down8” if standardized coverage was less than 3, “mid8” for coverage between 3 and 4.5, and “up8” for coverage exceeding 4.5. Among the S1 progeny, only “down7-up8”, “mid7-mid8” and “up7-down8” haplotypes, which respectively correspond to 0, 1 or 2 copies of tr8-7, were observed.

To analyze H08-1 dosage among MF93 S1 offspring, variants were jointly called using freebayes (version 1.3.4) with the 13,542 H08-1 SNP loci provided as targets. For each S1 offspring, read depth and H08-1 allele-specific read depth were aggregated across all loci, and the corresponding proportion of H08-1 specific alleles was used to determine H08-1 dosage in each offspring: dosage 0 was <5%, dosage 1 was 5-40%, dosage 2 was 40-60% and dosage 3 was ≥60%.

#### Trichome DNA extraction and genotyping

Trichomes were isolated from five Red Polenta leaves harvested from a single greenhouse-grown plant. Leaflets and stipules were manually removed. Petioles and petiolules were then cut into approximately 5 cm pieces, snap-frozen by immersion in a liquid nitrogen-chilled mortar, and gently scraped with a pre-chilled paintbrush and steel forceps to remove trichomes. Leaflets were then flash frozen in the same mortar, and longer trichomes were scraped from the midrib, veins and adaxial surface with steel forceps. Trichomes from all five leaves were ground to a fine powder under liquid nitrogen for approximately 15 minutes and carried forward for CTAB DNA extraction as described above. Trichome gDNA extracts were run on a 1% agarose gel to assess DNA concentration and quality. Approximately 20 ng of trichome DNA was amplified by combining 0.25 μM primer and 1X GoTaq Green Master Mix (Promega, USA) in a 50μl reaction volume. Reactions were subject to 3 minutes at 95°C, followed by 35 cycles of 30s 95°C, 30s at reaction-specific annealing temperatures, and 72°C for reaction-specific extension times. Reaction specific primer sequences, annealing temperatures and extension times are listed in Supplemental Table S2. All reactions were subjected to a 5:00 extension at 72°C. PCR products were purified by SeraMag SpeedBead cleanup and Sanger sequenced with the corresponding forward and reverse amplification primers by GeneWiz (South Plainfield, NJ). Sanger reads were aligned to the predicted DM1-3 amplicon sequence, and positions covering putative L1-specific alleles were identified in each read. At these positions, signal intensity of all four bases was extracted using the sangerseqR ^48^ and sangeranalyseR ^49^ packages in R (version 4.3.1), and the variant allele frequency calculated as the ratio of the channel intensity of the L1-specific allele to the total intensity of all channels at the position.

### Red Polenta genome assembly and assessment

#### Genome size estimation

Short reads were used to estimate the genome size of Red Polenta. Jellyfish version 2.2.7 ^50^ was used to count k-mers (k=31). The genome size and homozygous coverage level was then estimated from the 31-mer histogram output by Jellyfish using GenomeScope 2.0 ^51^.

#### De novo assembly

Primary contigs were assembled with hifiasm (version 0.15.5-r350) ^52^ with parameters ‘--primary --hom-cov 72 -t 48 --n-hap 4’ and all other parameters left at default settings. We proceeded with the processed unitig level assembly (output suffix “.p_utg.gfa”). Unitigs represent unique haplotypes that may be present one or more multiple times in a polyploid genome. Unitigs were classified according to dosage via read depth as previously described ^53^. Short reads were mapped to the unitigs with BWA mem (version 0.7.12-r1039) and long reads were mapped using minimap2 (version 2.17-r941) ^54^. For long read alignments, we filtered non-primary alignment with samtools version 1.9 (samtools view -q 1 -F 3840). Next, median read depth was calculated separately for each unitig, using the approach described above for the DM1-3 v6.1 assembly but with a sufficiently large window size to encompass each unitig. From examining the distribution of median read depths, we classified unitigs as haplotig [0 < x < 80], diplotig (80 < X < 135], triplotig (135 < x < 190], tetraplotig (190 < x < 245] or replotig (x > 245].

### Mutation detection

Short reads of Urgenta, Red Polenta and all protoplast regenerants were mapped to the Red Polenta unitigs with BWA mem (version 0.7.12-r1039). Alignments were preprocessed as described for DM1-3 v6.1 alignment processing, and raw variants were called with freebayes (version 1.3.4) with a minimum mapping quality of 40 and minimum base quality 20. Raw variant calls were filtered to retain biallelic SNP loci that were fixed for one of the two alleles in Urgenta with ≥3 supporting reads, and were not immediately adjacent to a run of three or more instances of the same nucleotide ^55^.

Layer-specific mutations were identified by comparing the genotypes of regenerants that carried tr8-7 with those that lacked tr8-7. To call a layer-specific mutation, we required that four of the six regenerants of one type exhibited a non-reference allele, while all six regenerants of the other type lacked the allele. We assumed that Urgenta carried the ancestral allele. Therefore, somatic mutation loci for which the Urgenta allele matched the L1-specific allele corresponded to loci in which a mutation arose and became fixed in the L2 and L3 layers of Red Polenta. In contrast, loci that exhibited the same allele in Urgenta and L2+L3-specific regenerants represent somatic mutations that arose and became fixed in the L1 layer of Red Polenta. To simplify this analysis, mutation counts are reported for haplotigs only, where heterozygosity is not expected. Additionally because they represent more reads, L2+L3-specific somatic mutations are more likely to lead to contig breaks during genome assembly than L1 mutations. This would result in a disproportionate number of L2+L3 specific somatic mutations near unitig edges, which present greater difficulties for mapping both short and long reads.

To define the assayable genome, we leveraged the slightly lower sequencing coverage of Urgenta short reads because it represented the limiting factor for mutation detection on haplotigs. Haplotig positions were considered assayed if more than three reads with mapping quality ≥ 40 and base quality ≥ 20 covered that position in the Urgenta reads.

Statistical tests for distribution of mutation types between layers were carried out as follows: layer-specific mutation counts were aggregated by transition or transversion, or alternatively, by the six canonical point mutation types, and differences between layers were compared using the prop.test() function in R (version 4.3.1). Counts of layer-specific mutations in each trinucleotide spectra were aggregated in R and passed to SigProfilerExtractor (version 1.1.4) ^27^.

## Data availability

Raw sequencing reads will be made available on NCBI Sequence Read Archive upon publication of this work.

## Acknowledgements

We thank Klaus J. Dehmer for providing genomic DNA of the IPK holding of Urgenta. This work was supported by the National Science Foundation Plant Genome Integrative Organismal Systems Grant 1444612 (Rapid and Targeted Introgression of Traits via Genome Elimination) to L.C. and I.M.H.

## Supplemental Information

**Supplemental File 1:** Supplemental Figures S1-S5

**Supplemental Table S1:** Sequencing information

**Supplemental Table S2:** Primers used in this study

