## Supplemental Figures for "Differential mutation accumulation in plant meristematic layers"

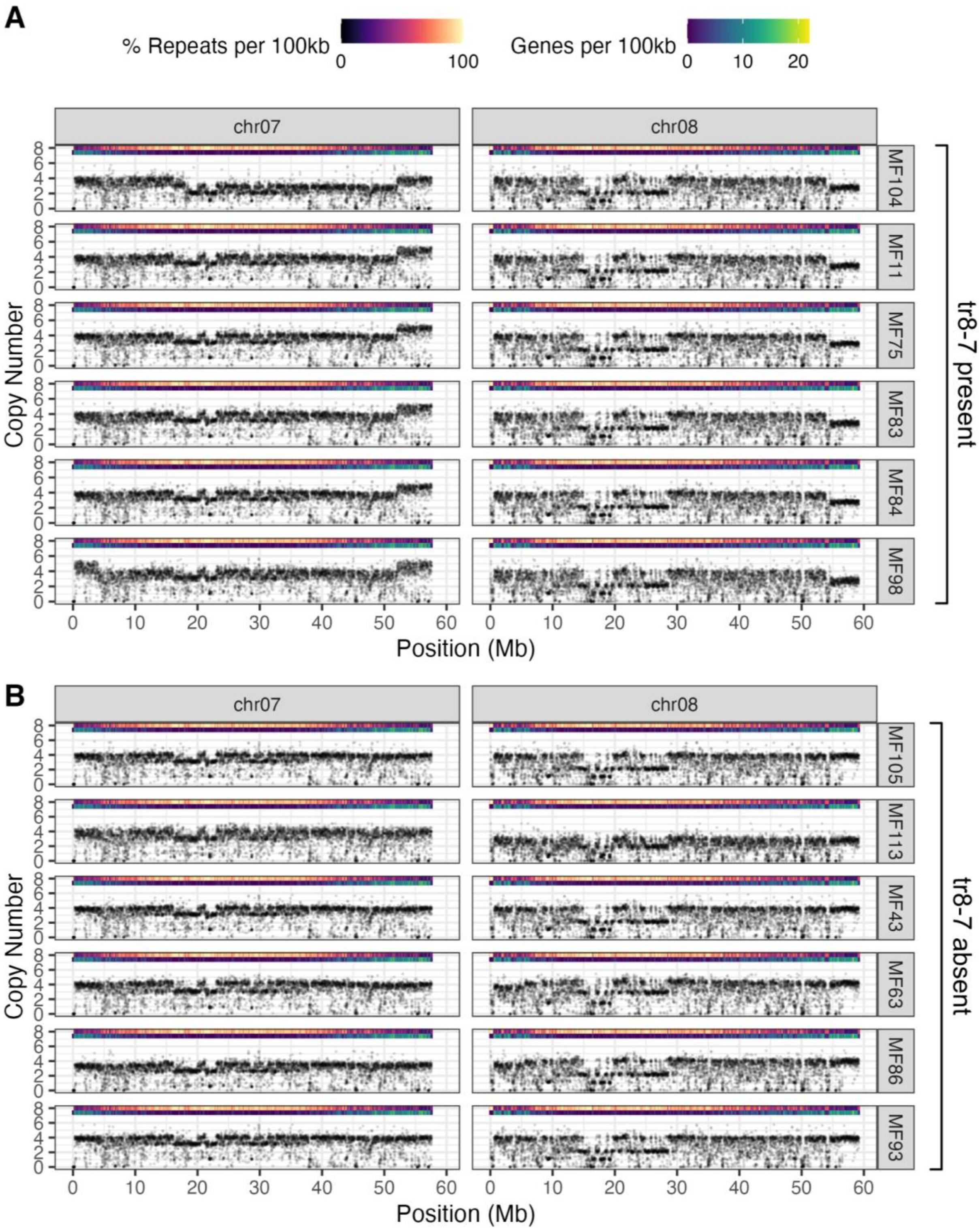


**Supplemental Figure S1.** Read depth analysis of regenerated leaf protoplasts of Red Polenta. Each data point corresponds to the median read depth of a non-overlapping 10kb window of the DM1-3 v6.1 reference genome, with only chromosomes 7 and 8 shown. **A)** Six regenerated lines exhibited tr8-7 after regeneration. **B)** Six regenerated lines lacked tr8-7 after regeneration.

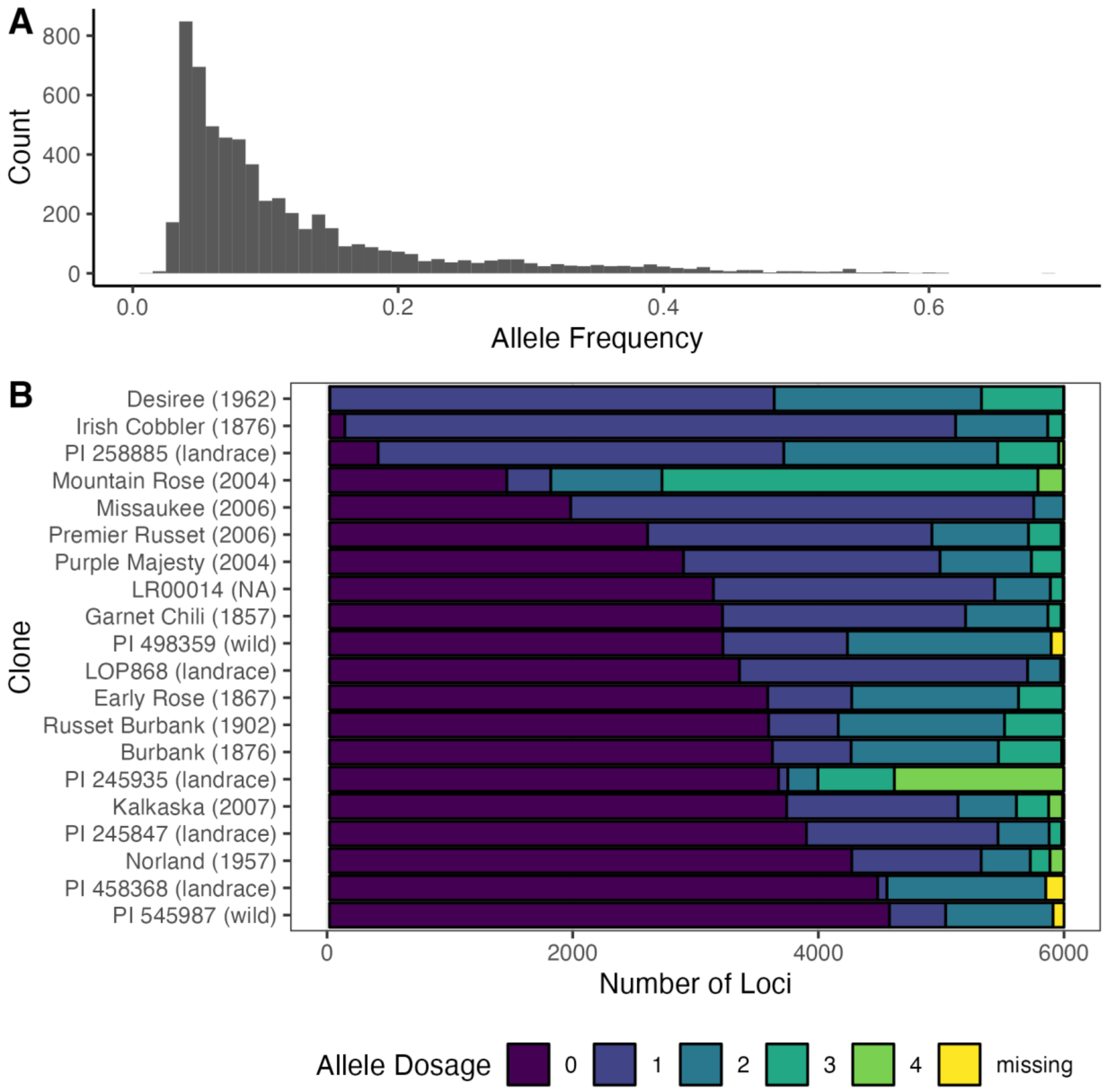


**Supplemental Figure S2. SNP alleles associated with tr8-7 absence in Red Polenta are common among potato varieties. A)** Frequency spectrum of approximately 6,000 SNP alleles associated with tr8-7 absence in Red Polenta among a potato diversity panel. Allele frequencies greater than zero indicate appearance of these alleles in other potato varieties. **B)** Bar plot illustrating the 20 individuals with the greatest extent of allele sharing among the diversity panel. Bar color indicates allele dosage. Allele sharing with landraces, wild species and varieties released before Urgenta in 1951 indicate reappearance of an ancestral haplotype. Pedigree release dates from [56].

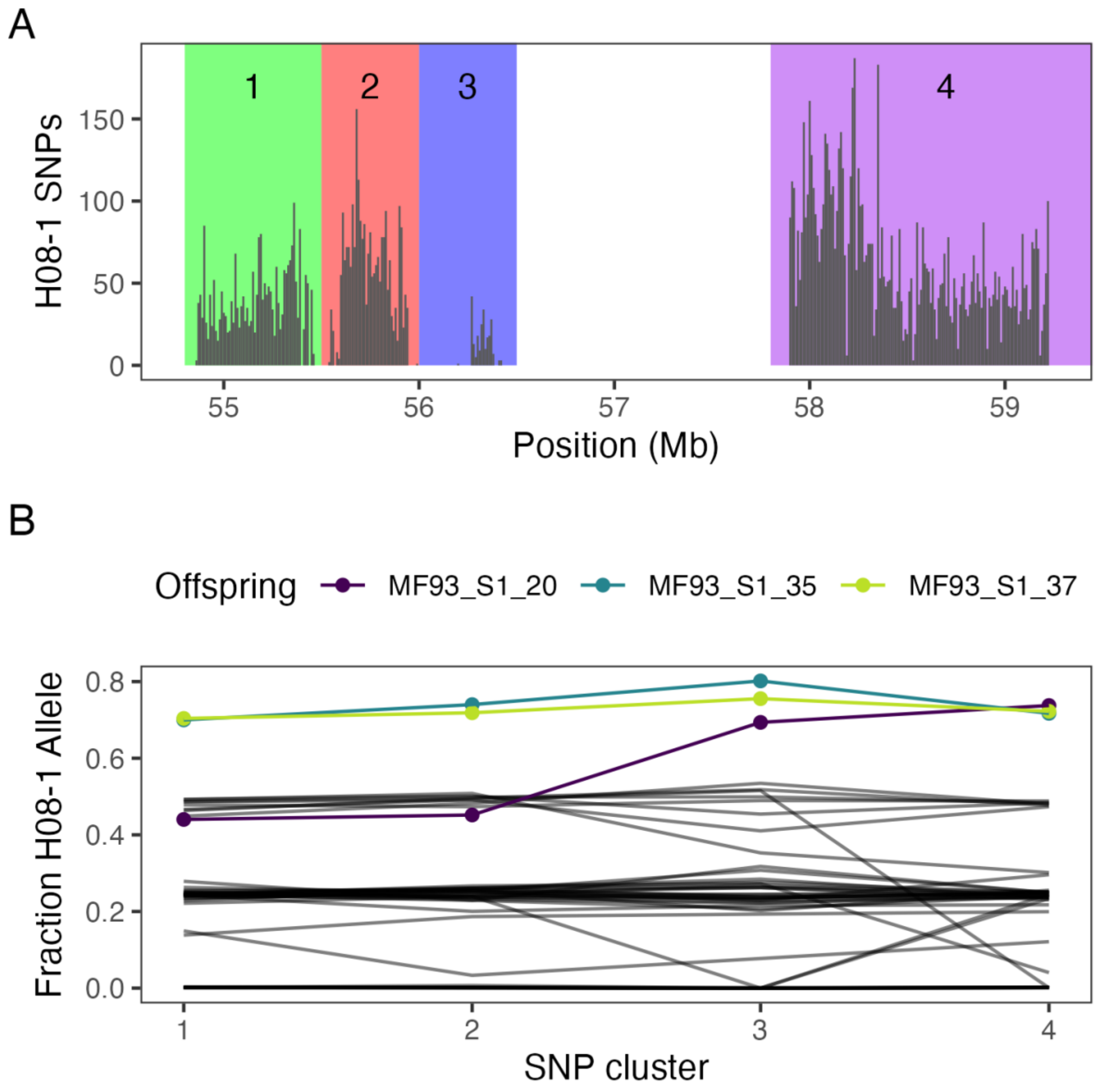


**Supplemental Figure S3. Evidence of double reduction in MF93 S1 progeny.** Three S1 progeny of MF93 exhibited H08-1 SNP allele dosage consistent with three copies of H08-1. **A)** Histogram of H08-1 SNP loci along the distal arm of chromosome 8. Bars correspond to the number of H08-1 SNP loci in non-overlapping 10kb windows. The region was divided into four clusters for a more detailed analysis of H08-1 dosage along chromosome 8. **B)** Fraction of H08-1 allele by cluster in MF93 S1 offspring. Individual offspring with H08-1 dosage exceeding 2 are shown as lines with unique colors; all other offspring are shown as black. Locations of crossovers involving the H08-1 chromosome are indicated by a change in the fraction of H08-1 allele between adjacent SNP clusters. Fractions exceeding 0.75 across all clusters are consistent with a crossover leading to double reduction between Cluster 1 and the centromere. A change in fraction from < 0.5 to approximately 0.75 is consistent with a crossover leading to double reduction between Cluster 2 and Cluster 3.

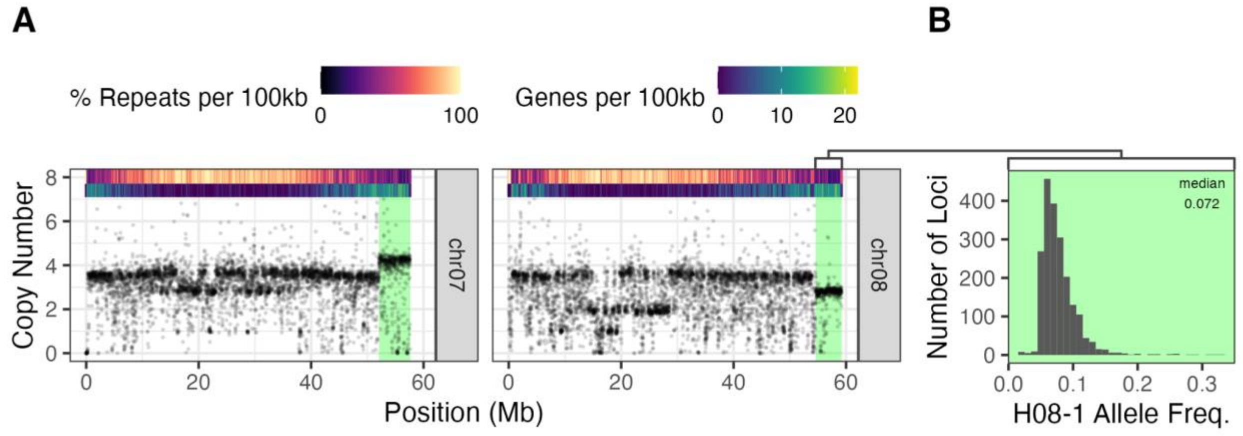


**Supplemental Figure S4.** A second Red Polenta holding exhibits the tr8-7 periclinal chimerism. **A)** Read depth of non-overlapping 10kb windows on chromosomes 7 and 8 reveals the tr8-7 pattern. **B)** Histogram of H08-1-specific allele frequencies, bin size 0.01.

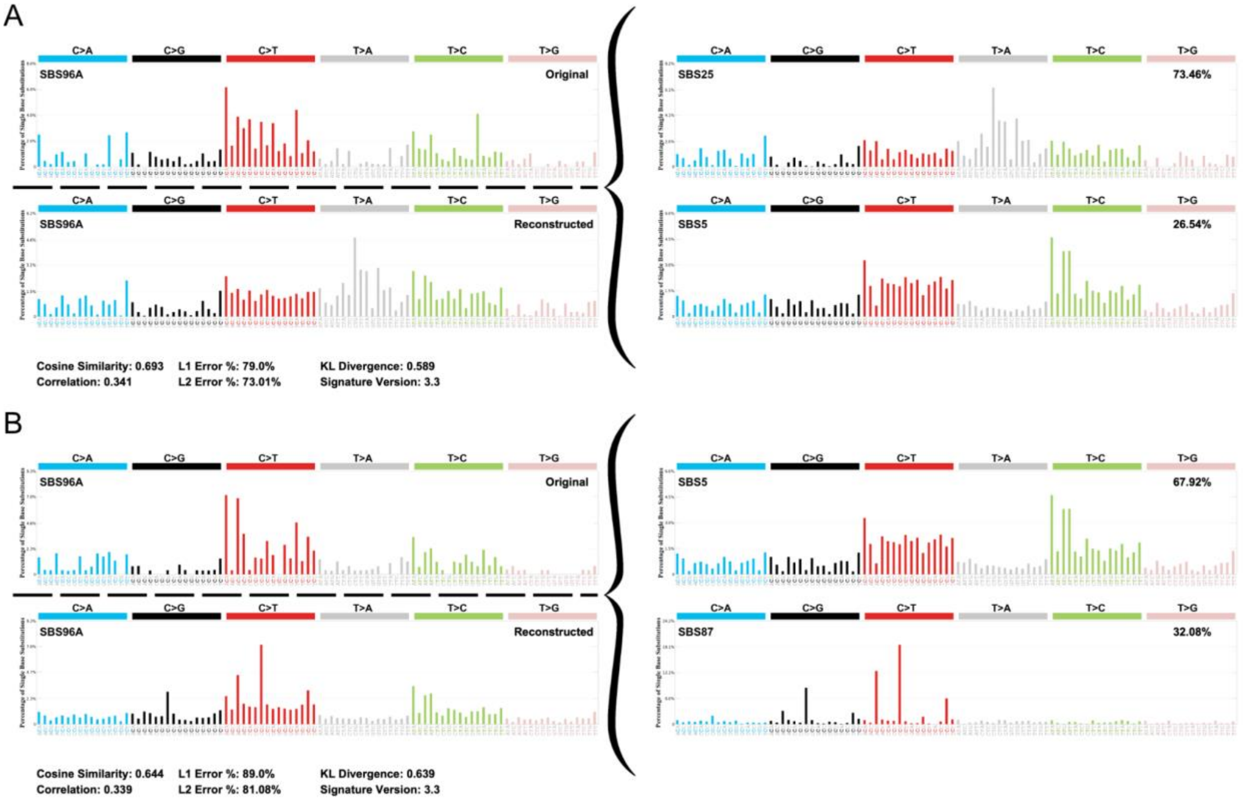


**Supplemental Figure S5**. Mutation signature profiling. The trinucleotide context of layer-specific mutations were extracted and compared to known trinucleotide mutational signatures of the COSMIC database. **A)** L1-specific mutations. **B)** L2/L3-specific mutations.
